# Malleability of the cortical hand map following a finger nerve block

**DOI:** 10.1101/2020.10.16.338640

**Authors:** Daan B. Wesselink, Zeena-Britt Sanders, Laura R. Edmondson, Harriet Dempsey-Jones, Paulina Kieliba, Sanne Kikkert, Andreas C. Themistocleous, Uzay Emir, Jörn Diedrichsen, Hannes P. Saal, Tamar R. Makin

**Affiliations:** Institute of Cognitive Neuroscience, University College London (UK); Wellcome Centre for Integrative Neuroimaging, University of Oxford (UK); Department of Neurobiology, Harvard Medical School (USA); Active Touch Laboratory, Department of Psychology, The University of Sheffield (UK); Nuffield Department of Clinical Neurosciences, University of Oxford, Oxford, UK & Brain Function Research Group, University of the Witwatersrand, Johannesburg, South Africa; Brain and Mind Institute, University of Western Ontario (CA); Wellcome Centre for Human Neuroimaging, University College London (UK)

**Keywords:** Plasticity, primary somatosensory cortex, remapping, reorganisation, fMRI, deprivation, deafferentation

## Abstract

Individual fingers in the primary somatosensory cortex (S1) are known to be represented separately and adjacently, forming a cortical hand map. Electrophysiological studies in monkeys show that finger amputation triggers increased selectivity to the neighbouring fingers within the deprived S1, causing local reorganisation. Neuroimaging research in humans, however, shows persistent S1 finger representation of the missing hand, even decades after amputation. We aimed to resolve these apparently contrasting evidence by examining finger representation in humans following pharmacological ‘amputation’ using single-finger nerve block and 7T neuroimaging. We hypothesised that beneath the apparent selectivity of individual fingers in the hand map, peripheral and central processing is distributed across fingers. If each finger contributes to the cortical representation of the others, then localised input loss will weaken finger representation across the hand map. For the same reason, the non-blocked fingers will stabilise the blocked finger’s representation, resulting in persistent representation of the blocked finger. Using univariate selectivity profiling, we replicated the electrophysiological findings of local S1 reorganisation. However, more comprehensive analyses confirmed that local blocking reduced representation of all fingers across the entire hand area. Importantly, multivariate analysis demonstrated that despite input loss, representation of the blocked finger remained persistent and distinct from the unblocked fingers. Computational modelling suggested that the observed findings are driven by distributed processing underlying the topographic map, combined with homeostatic mechanisms. Our findings suggest that the long-standing depiction of the somatosensory hand map is misleading. As such, accounts for map reorganisation, e.g. following amputation, need to be reconsidered.

## Introduction

Representation in the primary somatosensory cortex (S1) has long been conceptualised as a somatotopic gradient, i.e. neighbouring sub-regions are selective in their responses to neighbouring parts of the body. The primate hand area is a well-established model for such somatotopic organisation. Traditionally, the hand area has been divided into discrete clusters, each selective to a single finger, lined up mediolaterally – hereafter referred to as the *hand map* (Kaas et al. 1979, Sanchez-Panchuelo et al. 2010, Martuzzi et al. 2014). The hand map has been useful for studying adult plasticity, as changes to the spatial organisation of the finger clusters are relatively easy to monitor (Buonomano and Merzenich 1998). Changes in the boundaries of the finger representation have been largely attributed to Hebbian plasticity, i.e. positive feedback mechanisms (Allard et al. 1991, Wang et al. 1995), but it is important to note that homeostasis, i.e. negative feedback mechanisms are also important in regulating activity-dependent plasticity. These opposing forces can both work selectively on single synapses, but also at the network level (Turrigiano and Nelson 2004).

Foundational literature on somatosensory plasticity, mainly in animal models, shows that the boundaries between fingers within the hand map are profoundly altered following localised input loss. These studies demonstrated that the deprived cortical territory becomes activated by inputs from the cortically-neighbouring body parts (Merzenich et al. 1983a, Pons et al. 1991, Churchill et al. 1998). For instance, following amputation of the middle finger (D3) in new-world monkeys, Merzenich et al. (1984) described an invasion of the neighbouring fingers (D2 and D4) into the deprived cortical territory. Multiple processes have been offered to drive such dramatic remapping, including anatomical changes (Rasmusson 1982). Yet, consensus in the field now favours the unmasking of previously silent inputs as a key process (Kaas 1991, Weinberger 1995, Feldman and Brecht 2005). These findings have been seminal in revealing the ability of primary cortical areas to be plastic following changes in experience occurring in adulthood.

However, the results described above are in contrast with recent evidence in humans documenting the limited effect of amputation on functional organisation of hand representation. We (Kikkert et al. 2016, Wesselink et al. 2019) and others (Bruurmijn et al. 2017, Serino et al. 2017) have shown that even decades after amputation of their hand, human amputees continue to represent the fingers of their missing hand, with similar canonical organising principles as in typical S1 hand representation. Using various functional MRI measurement techniques, these studies indicate the representational features of the hand are preserved within the sensorimotor system despite input loss. For example, phantom hand movements evoke activity patterns in S1 that are largely indistinguishable from normal hand movement (Wesselink et al. 2019). The classic and more recent human studies differ in experimental species, acquisition techniques and manner of stimulation (mainly passive stimulation versus active movement), which could explain some differences in the main findings. Yet ultimately, the persistence of hand representation in humans does not match the idea that the deprived cortex is rapidly overwritten.

With regard to the organising principles underlying the S1 hand map, several more recent studies provided converging evidence that diverge from the classical view of somatotopic single-finger representation. First, Ejaz et al. (2015) have shown that the statistics of finger (co-)use measured in daily life provides a better explanation of S1 hand representation than simple somatotopy. Specifically, using representational similarity analysis (see Methods), they showed that cortical activity patterns for different fingers overlap considerably and form a more complex structure than suggested by somatotopy alone (see also Besle et al. 2014, Kuehn and Pleger 2020). Secondly, Manfredi et al. (2012) have demonstrated that even passive stimulation of a part of the hand triggers extensive ripple effects across the skin. Because these vibrations activate mechanoreceptors on other fingers, localised tactile stimulation elicits widely distributed peripheral responses, with distinct spatiotemporal patterns (Shao et al. 2016, Shao et al. 2020). Both of these results suggest that shared somatosensory processing of inputs from multiple skin surfaces across the hand might be more prevalent than previously anticipated. Therefore, inter-finger representation might play a more important role in shaping central finger representations. As such, it is worthwhile to re-examine the seminal findings on somatosensory plasticity and the role of cortical neighbourship. If tactile processing is more distributed and the different finger areas more inter-connected than typically thought, then any local input change should lead to far-reaching changes across the cortical hand map. Although subtle global changes to the hand map, i.e. an altered selectivity profile outside the deprived cortex, were already observed by Merzenich et al. (1983b), the vast majority of studies have focused on characterising local reorganisation within the deprived cortex, leaving more global plasticity relatively unexplored.

In this study, we mimicked the rapid cessation of somatosensory input typically associated with amputation, using a pharmacological nerve block, allowing us to longitudinally characterise deprivation-trigger changes to the human S1 hand map. Healthy volunteers underwent two sessions of hand mapping experiments using ultra-high-field fMRI (1mm resolution), one while their index finger was locally blocked using a pharmacological anaesthetic agent (block session) and another without a block (baseline session; see Figure 1A for experimental timeline). To modulate the level of afferent and efferent input, which was different between human and animal amputation experiments, we probed S1 with both passive (tactile stimulation) and active (finger tapping) stimulation (see Sanders et al. (2019) for a comparison between active/passive conditions on S1 hand representation at baseline). We also acquired resting-state fMRI and magnetic resonance spectroscopy, both of which can provide task-independent markers of plasticity across the hand map. We tested the idea that persistent representation can result from more overlapping finger representations of the fingers than typically considered. We hypothesised that due to peripheral and central processing being shared across fingers, the blocked finger’s missing activity can be reinstated by the other fingers. With regards to classical accounts of deprivation-triggered remapping, we predicted that if fingers are more overlapping across the hand map to begin with, then simply by removing the dominant input from a given finger cluster we will expose the neighbouring fingers representation. But importantly, since this wouldn’t necessitate any synaptic change, deprivation should not drive increased activity for the neighbouring non-blocked fingers. Instead, we hypothesised that due to the shared inter-finger representation, reduced input to one finger should decrease activity for the other (non-blocked) fingers.

**Figure 1:**
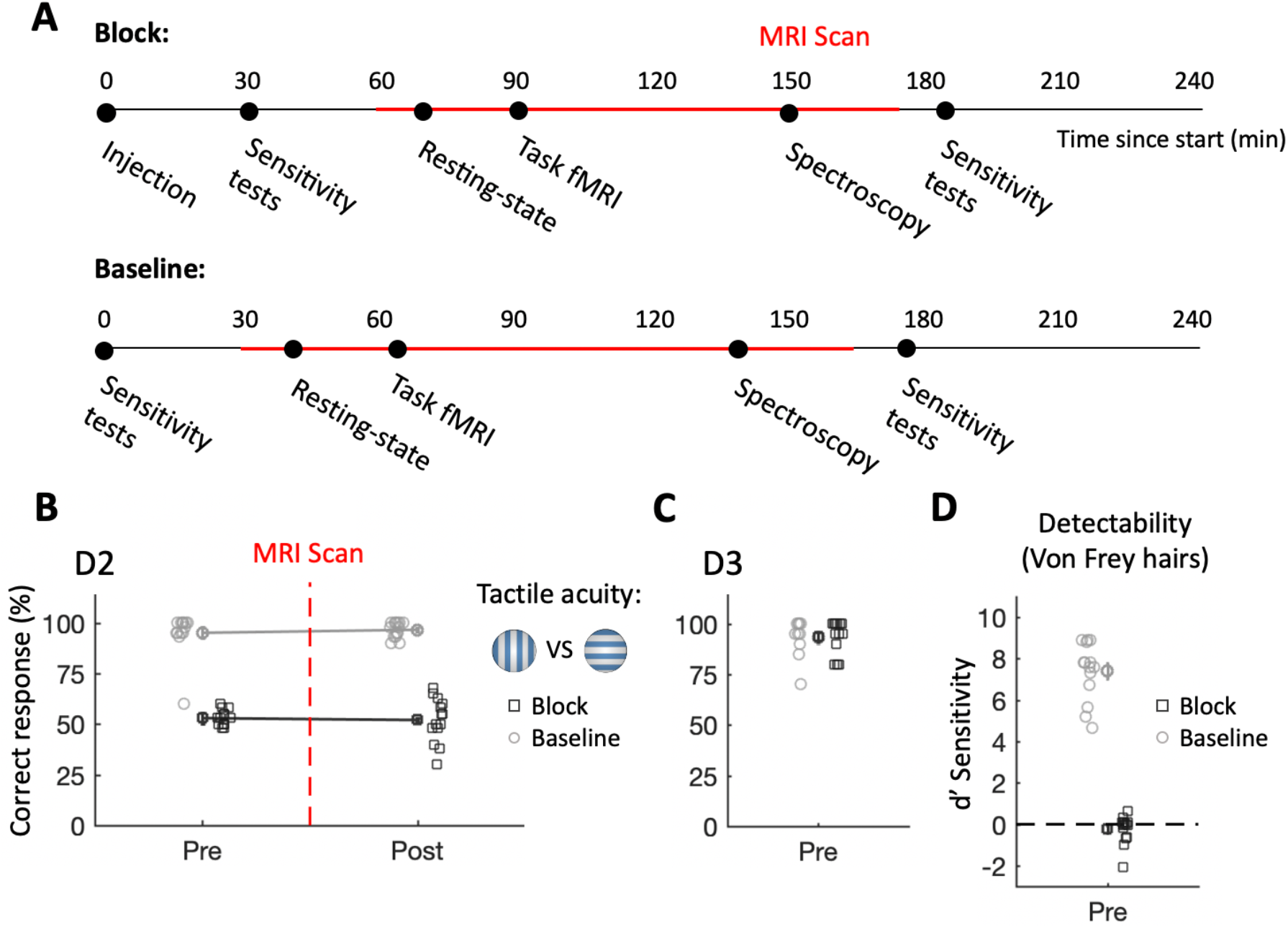
Successful administration of a local nerve block. **A)** Experimental timeline. Dots indicate the approximate start of each study component. Time spent in the MRI scanner is marked in red. Both sessions were identical except for the injection that was only administered in the block session and the independent localiser task that was only performed in the baseline session. **B)** Tactile acuity, assessed using a grating orientation judgement task, is abolished for the index finger (D2) following a local nerve block. During the baseline session, participants had near-ceiling performance. Performance remained stable throughout each experimental session. The (categorical) x-axis is jittered for visualisation purposes. **C)** Tactile acuity was unaffected in the non-deprived and neighbouring middle finger (D3), confirming that our intervention was localised to D2. **D)** Detectability of light touch in finger D2 drops to chance level following nerve block.

## Results

The use of active and passive tasks was originally aimed at resolving some possible causes of methodological divergence in human and animal studies. However, given overall consistency in the results between the active and passive conditions, the analyses reported below focus on the passive condition; parallel results from the active condition are reported in the supplementary section. We highlight instances where results diverged between the two paradigms.

### Local nerve block induces abolishment of tactile acuity and local activity

We first established that perception for the index finger (D2) was effectively obliterated in the block session using a tactile grating orientation judgement test. All participants showed near-ceiling performance in the baseline session (95%, see Figure 1B), whereas performance dropped to chance level after the nerve block (53%). Performance remained poor after the end of the scan, some three hours after the block was administered (main effect of session: χ^2^(1) = 839.97, p < .001; time x session interaction: χ^2^(2) = 0.32, p = .570). Before the scan (and after the nerve block in the block session), we also tested for reductions in perception of light touch, the slowest somatosensory modality to be abated by a nerve block (Raymond and Gissen 1987), using Von Frey hairs (Figure 1D). Participants’ sensitivity to detect light touch was reduced from nearly perfect to chance level after the nerve block (χ^2^(1) = 324.73, p < .001). We also tested the acuity of the neighbouring middle finger, to ensure that the effects of the nerve block had not spread to other fingers. We noted near-ceiling grating orientation performance in both sessions prior to the scan (and after the block in the block session; baseline: 92%; block: 93%; main effect of session: χ^2^(1) = 0.13, p = .723; Figure 1C). At the end of each session, we directly assessed the participants’ perceptual thresholds for the middle (D3) and ring finger (D4) using gratings (see Methods for details). There was no significant main effect of session (χ^2^(1) = 1.03, p = .311), nor a finger x session interaction (χ^2^(2) = 0.37, p = .541), further supporting that the effect of the nerve block had not spread to the non-target fingers.

We next demonstrated that the nerve block was physiologically effective in diminishing D2 activity in its respective S1 area. To characterise finger representation based on univariate activity, we first identified five “finger clusters” (C1-5), each consisting of only the voxels showing strong selectivity to one finger over the other four. Clusters were identified in each participant’s hand area by means of a standard travelling wave analysis performed on an independent localiser map (Figure 2A, see Methods). Within each cluster, mean finger-specific activity levels were obtained from the randomised block design task collected during both baseline and block sessions (Figure 2B). As expected, after the nerve block, passive stimulation of D2 no longer induced activity in its corresponding finger cluster (C2) relative to rest (μ=−0.04; t_(14)_=−0.10, p=.919, Bayes’ Factor=.266; decrease versus baseline session; t_(14)_=−4.17, p=.001). Active D2 stimulation still elicited positive activity in C2 (μ=2.13; t_(14)_=4.40, p=.001; see Figure S1A), presumably due to additional afferent (e.g. proprioceptive inputs from the arm muscles) and efferent signals (from the motor system). Note however that in the active condition, the activity was also significantly reduced in the block condition relative to baseline (t_(14)_=−2.22, p=.044).

**Figure 2:**
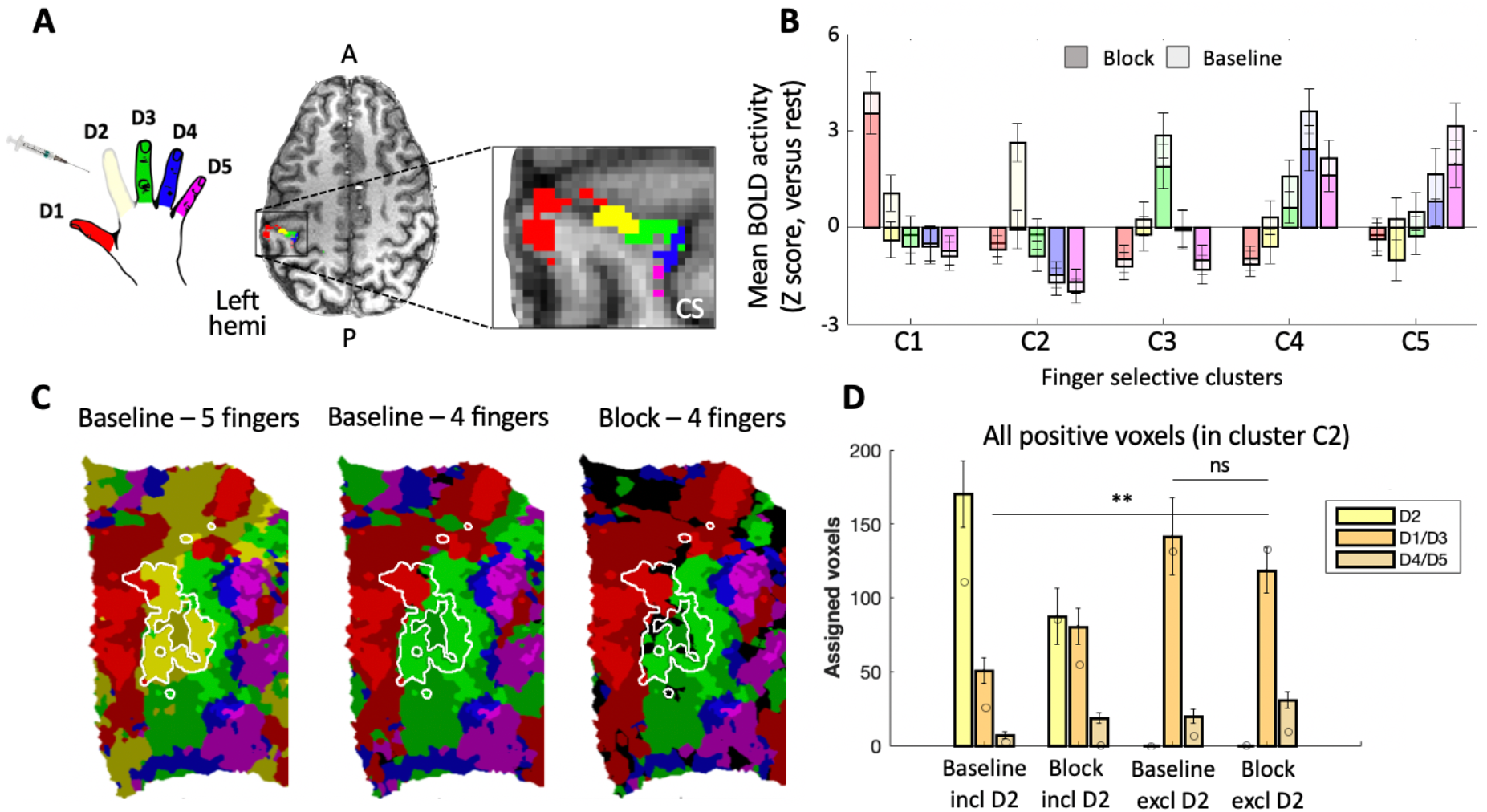
Single finger nerve block attenuates D2 activity but does not cause true reorganisation – evidence from univariate analysis. **A)** An example of the five finger clusters that were localised in S1 for each individual participant. **B)** Activity levels for each of the five fingers (colours as in A) across the five finger clusters in the baseline (light colours) and block (dark colours) sessions; Group results for passive stimulation. In the baseline session, activity profiles across fingers reflect classical finger topography. In the block session, activity for finger D2 is abolished in cluster D2, but also decreases for other target fingers in their respective clusters. Error bars indicate standard error of the mean. **C)** Winner-takes-all assignment of voxels to fingers, presented on a flattened surface for an example participant, for positive (above-zero) activity only (colours as in A). The white outline shows the D2-selective cluster, as identified independently. Compared to measuring 5 fingers in the baseline sessions (left), not measuring finger D2 reveals similar “invasion” of the neighbouring fingers (D1 – red and D3 – green) into the D2 territory (white outline), irrespective of a nerve block, resulting in similar “remapping” in baseline as found in the block map. Bright colours indicate voxels within the independently-localised finger clusters; shaded colours are activated voxels (Z>0) outside the independently-determined finger clusters; black indicates below-zero activity. **D)** Quantification of the remapping effect demonstrated in C; Group results. Voxels within the D2 cluster were assigned to either one of 5 fingers (including D2) or one of 4 fingers (excluding D2) using a winner-takes-all approach. The number of voxels assigned to D2 (when included), neighbouring fingers (D1/D3) and non-neighbouring fingers (D4/D5) were calculated per participant and averaged. Note that due to the independence of the tested data from the cluster definition, not all voxels may be assigned to D2 in the ‘baseline’ session. Increased representation of the neighbouring (D1/D3) digits in the D2 cluster was not significantly different between the baseline and block sessions. Error bars indicate standard error. Circles indicate the values for the example participant that was showcased in C. Abbreviations: A: Anterior; P: Posterior; Left hemi: Left hemisphere; ns: non-significant; **: p<.001.

Activity elicited by (passive) stimulation of D2 was also reduced in the other four finger-selective clusters compared to baseline (F_(1,112)_=9.90,p=.002), with no significant interaction across clusters (F_(3,112)_=0.48, p=.700). Thus, the nerve block successfully attenuated D2 univariate activity across the hand area.

### Local ‘remapping’ might reflect analysis choices rather than neural plasticity

Classical electrophysiological studies involving amputation have reported an overrepresentation of neighbouring fingers in the deprived territory. In order to use methods similar to those studies, we first calculated winner-takes-all maps (Figure 2C), rather than using the full activity patterns. In these maps, all voxels showing positive (above zero) activity to one of the fingers were assigned to the finger with the highest activity level (relative to baseline). Similar findings were observed when activity was not thresholded, such that negative BOLD activity was also considered (see Figure S2).

The finger preference of the voxels in cluster C2 (defined based on the independent travelling-wave localiser map) was measured by counting the number of voxels assigned to the neighbouring fingers. We first repeated the classical analysis - ignoring the D2 condition in the winner-takes-all map of the block session –– a necessity in animal studies involving amputation –– but not in the baseline session. When comparing the number of assigned voxels across sessions, we find a significant increase in the number of voxels that show a preference for D1 or D3 after a nerve block, compared to the baseline session (t_(14)_=5.10, p<.001; see Figure 2C for an example map and Figure 2D for group comparisons). This is consistent with previous accounts of local remapping, caused by over-representation of neighbouring fingers. However, a key advantage to our experimental model is the possibility to stimulate D2 after deprivation. When D2 responses are included in the winner-takes-all analysis, the remapping of neighbouring fingers D1 and D3 in C2 is merely trending (t(14)=2.13, p=.051). Furthermore, simply ignoring D2 in the baseline data already gives the impression of shifted finger boundaries (4 fingers winner-takes-all relative to the 5 fingers comparison, both at baseline; t(14)=4.84, p<.001; Figure 2D). Conversely, when D2 was ignored in both sessions (i.e. excluded from the winner-takes-all analysis), we found no significant effect of the nerve block on neighbouring fingers ‘remapping’ in the deprived cluster (t(14)=−1.12, p=.280). This suggests that the previously observed findings might have been confounded by methodological restrictions. In other words, remapping or other mechanisms of plasticity may not be necessary for explaining a change in the winner-takes-all maps.

The conclusion that remapping of the cortical neighbours into the deprived cortex may not have occurred in our study was further supported by other analysis, focusing on mean activity within the finger clusters. Mean activity in C2 for fingers D1 and D3 did not significantly increase in the passive condition (t(14)=−1.09, p=.304; Figure 2B), and had, in fact, decreased in the active condition (t(14)=−2.51, p=.025). Together, these results suggest that the changes in the hand map came about by uncovering pre-existing co-activation of voxels by fingers other than the ‘winning’ (target) finger.

### Global loss of selectivity in the hand region

Given the hypothesised overlap in finger representation, we also investigated the possibility for more global changes in the hand representation, i.e. outside the deprived area. We first examined whether responses within the finger clusters beyond C2 were affected by the nerve block by assessing univariate finger selectivity: the activity for each cluster’s target finger minus the mean of the three non-target fingers, excluding D2 (see the Supplementary Section for the same analysis while including D2). Finger selectivity is considered a hallmark of sensory cortical organisation, particularly in S1. With passive stimulation, we identified a significant decrease in selectivity across all (non-blocked) finger clusters (mean change: −21.7%; F_(1,112)_=8.14, p=.005), and no session x cluster interaction (F_(3,112)_=0.31, p=.818). This result provides clear evidence for larger-scale changes in activity across the entire hand map, which have not previously been considered. Indeed, when examining Figure 2B it is apparent that the non-target fingers are not equally suppressed within a given cluster, but vary in a systematic fashion, in line with topography. That is, neighbouring fingers that were more strongly activated at baseline also show greater suppression.

To investigate the global selectivity reduction in more detail, we conducted representational similarity analysis (RSA; see Methods). RSA produces a canonical S1 representational structure for ‘normal’ hands based on inter-finger similarity patterns, and can therefore be used to assess variations in inter-finger dissimilarity. While extensively previously used to characterise the fingers representational structure across the entire hand area, here we used the same approach to identify representational similarity within each finger-selective cluster. For each cluster, the dissimilarity between all individual finger patterns, as well as rest, was compared in each session. For visualisation purposes, the representation patterns were projected onto a two-dimensional plane using multi-dimensional scaling (Figure 3A), such that distances in the projection reflect dissimilarity between conditions (for representational dissimilarity matrices, see Figure S3). Given that we used cross-validated distances (see methods), systematically positive differences imply statistically reliable differences between patterns.

**Figure 3:**
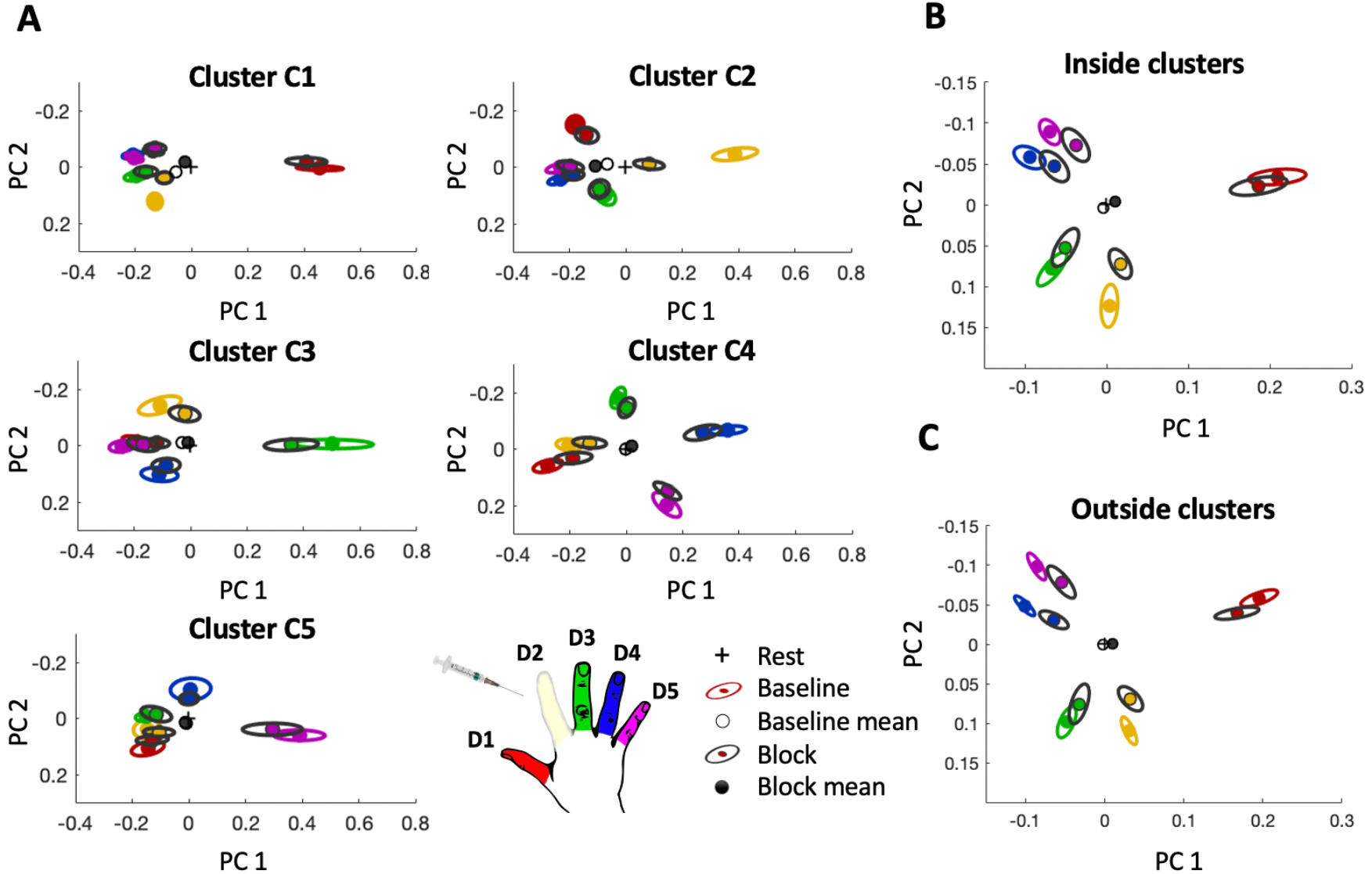
Finger representation is distributed throughout the hand area and globally affected by a local nerve block. A) In each finger cluster (C1-5), both target and non-target fingers show differential activity patterns from each other and from rest. Fingers are coloured according to the hand-shaped legend; the surrounding ellipses (indicating between-subject standard error) are either coloured (baseline session) or black (block session). The inter-finger representational structure (comprised of all five fingers) is relatively consistent across sessions. B) When all clusters are examined simultaneously, the prototypical hand representation shrinks following the nerve block, but does not change shape. C) Hand representation *excluding the finger-selective clusters C1-C5* strongly resembles that of the highly selective voxels, suggesting hand information is distributed throughout the S1 hand area.

We first consider the inter-finger representational structure in each of the finger-selective clusters in the baseline session. Selectivity of each of the finger clusters as defined above can be inferred with RSA by the target finger showing highest dissimilarity from rest, relative to the non-target fingers (Figure 3A; colours). The non-target fingers in each of the five clusters, however, can also be distinguished from each other (mean dissimilarity between non-target finger pairs per cluster; t’s_(12-14)_>5.90, p’s<.001; one-sample t-tests). In three out of five clusters, the dissimilarity distance rank between non-target fingers (e.g. D3 is closer to the D4 than to D5) was preserved (average rank-order correlation with the canonical hand pattern, i.e. the pattern from the entire hand area in the baseline session: C1: rho=.64, p=.038; C2: rho=.55, p=.024; C3: rho=.26, p=.083; C4: rho=.46, p=.122; and C5: rho=.51, p=.008). This suggests that at baseline, finger information is highly distributed, such that even the most finger-selective clusters contain whole-hand topographic information. Therefore, in order to study the impact of the nerve block on S1, the entire hand area should be examined.

We next interrogated the impact of the D2 nerve block on the representational structure in an anatomically defined region around the “hand knob” spanning the entire S1 hand area (see Methods; Further analysis on the impact of the nerve block on the representational structure in the finger-selective clusters is elaborated below). To avoid overlap with the observations presented above we excluded all voxels with pronounced selectivity for a single finger, i.e. voxels inside finger clusters. Akin to the reduction in finger selectivity in Figure 2, inter-finger dissimilarity due to passive stimulation was significantly reduced in the extended hand area across the 4 non-blocked fingers following the nerve block (−29%, t_(13)_=3.00, p=.010). Neighbouring fingers (D1 and D3) were not affected to a significantly different extent than non-neighbours (D4 and D5; finger neighbourhood x session interaction: F_(1,52)=._12, p=.731). Similar results were also observed when only the activity underlying the finger-selective clusters was examined (−26%, t_(13)_=2.54, p=.025; Figure 3B). As can be seen by comparing Figures 3B and C, the representational structure in the most finger-selective voxels and the non-selective voxels comprising the anatomical hand area is highly similar (mean within-subject correlation: baseline session: r=0.88; block session: r=0.89).

There was a similar drop in dissimilarity during the active task (inside clusters: −19%, t_(14)_=3.45, p=.004; outside clusters: −21%, t_(14)_=4.40, p=.001; Figure S1), also without a significant finger neighbourhood x session interaction (F_(1,56)_=.20, p=.660). Importantly, the measured drop in dissimilarity could not, to the best of our knowledge, be attributed to inter-session differences that are unrelated to the nerve block, e.g. task performance or brain state. We determined that there were no significant differences between sessions in press-force during (active) task performance (t_(14)_=−0.81, p=.433) or perceptual detection of an odd-ball trial during (passive) task performance (t_(14)_=0.34, p=.737); the residual error in the GLM (t_(14)_=0.86, p=.405); or the signal-variance in S1 in a separate resting-state scan (t_(12)_=0.40, p=.700, see methods). Collectively, the above findings provide converging evidence that individuated finger representation became less distinct across the hand area following D2 block. Although the deprivation of input was local, i.e. affecting only one finger, finger information is distributed, leading to widespread effects on the representation of the non-blocked fingers across the entire hand map.

### Persistent representation of the blocked finger is found outside the C2 cluster

From our analysis so far, it becomes apparent that the reduction of D2 activity via nerve block affected non-blocked fingers representation across the hand area. We next explored the idea that, due to distributed and overlapping finger representation, activity associated with one finger may inform the representation of another finger, or could even reinstate it in the case of missing input (e.g. D2 representation in the block session). Going back to the (passive) cluster-specific analysis in Figure 3A, the blocked D2 representation is significantly different from the rest condition in all clusters (t_(13)_’s>3.87, p’s<.002). Looking at the specific effects of the D2 block, it appears that while D2 dissimilarity drops sharply compared to rest in cluster C2, in the other clusters it is more stable. Statistically, we identified a significant dissimilarity drop in clusters C1 (t_(13)_=2.50, p=.027) and C4 (t_(13)_=2.43, p=.031), but not in clusters C3 (t_(13)_=1.09, p=.297) and C5 (t_(13)_=1.21, p=.247). Importantly, as exemplified in the hand-area analysis detailed above, this dissimilarity drop is not specific to D2. When each cluster’s four non-target finger are examined compared to rest, no cluster shows a significant session x finger interaction (0.11<F_(3,104)_<0.81; p’s>.489), yet every cluster showed a main effect of session (F_(1,104)_’s>7.73; p’s<.008). In other words, while we see a general decrease in finger-specific information, this decrease seems relatively stable across the non-blocked fingers. The global dissimilarity reduction across the non-blocked fingers was also true when the fingers were grouped as neighbours and non-neighbours to the target finger (effect of session: F_(1,108)_’s>6.90, p’s < .010; session x neighbourhood interaction: 0.00<F_(1,108)_<1.64, p’s > .204). These findings stand in stark contrast to the idea of increased selectivity of the cortical neighbours due to local cortical remapping.

To test whether the blocked D2 sustained greater reduction in representation relative to the other fingers, we compared the entire hand area (excluding the finger-selective clusters; Figure 3C). We again do not find significant evidence for a greater collapse in dissimilarity from rest for any of the fingers (including all 5 fingers; effect of session: F_(1,130)_=5.99, p = .002; session x finger interaction: F_(4,130)_=0.07, p = .991). Therefore, the reduction in information about D2 as a consequence of the nerve block was, with the exception of cluster C2, not strikingly different from that about the non-blocked fingers. In other words, the localised nerve block triggered a relatively homogenous reduction in dissimilarity across the entire hand.

In the active condition, no cluster showed a significant session x finger interaction (0.20<F_(3,112)_<1.12, p’s>.344). Only C4 showed a significant main effect of session (C4: F_(1,112)_=18.46, p<.001; other clusters: 0.15<F_(1,112)_<3.07, p’s>.083). Active results from the entire hand area (excluding the finger-selective clusters) are in line with that of the passive condition (including all 5 fingers; effect of session: F(1,140)=10.7, p=.001; session x finger interaction: F(4,140)=0.21, p =.930). Together, this suggests that, despite the reported reduction in D2 activity across the hand map, the finger’s position within the canonical hand representational structure is preserved.

### Homeostatic mechanisms can support global representational stability

To determine which factors may have contributed to the observed large-scale effects despite local peripheral deprivation, we used a computational model that simulates S1 hand representation based on peripheral inputs (see Methods for details). In short, this model makes explicit that passive finger stimulation causes ripples in the skin that can activate mechanoreceptors beyond the immediate stimulation site (Saal et al. 2017). Based on the typical distribution of slowly and rapidly adapting (SA1, RA), and Pacinian corpuscle (PC) cutaneous mechanoreceptors, profiles of afferent responses to modelled stimuli can be generated (Figure 4A). The cortex is modelled with five units (representing the five cortical finger-selective clusters) that receive input from the periphery and are also connected laterally (Figure 4B). Indeed, including broad inter-connectivity between different cortical patches allowed the model to encapsulate the canonical two-dimensional hand representation in S1 as found using RSA in this and previous studies (Ejaz et al. 2015, Mehring et al. 2019, Kieliba et al. 2020). Importantly, the model allowed us to disambiguate which observed activity changes in the anaesthetic condition can be attributed to the loss of peripheral input alone, and which require additional central plasticity mechanisms.

**Figure 4:**
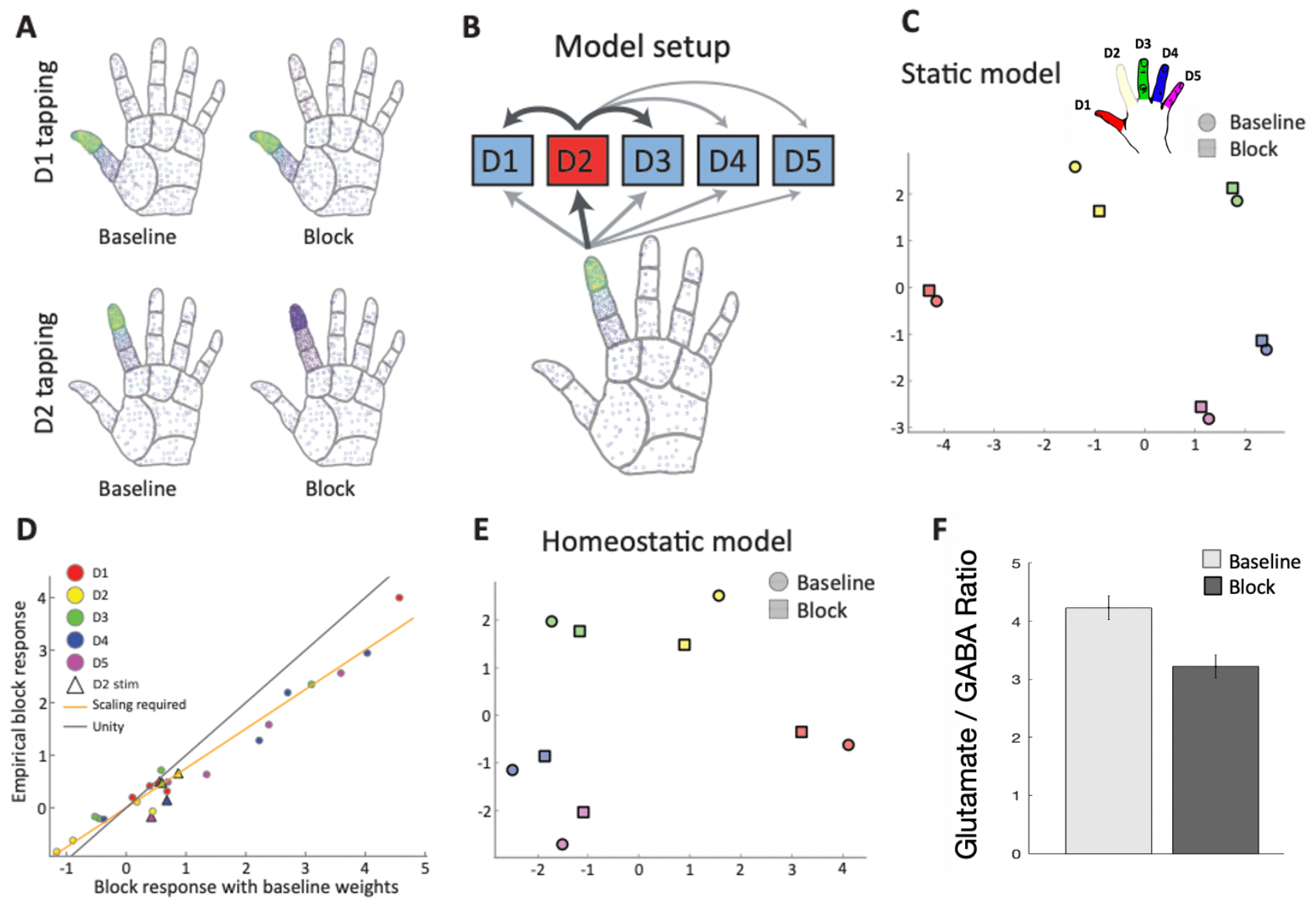
A computational model implementing a combination of local peripheral block and homeostatic mechanisms can account for the observed results. **A)** Simulated peripheral responses to mechanical finger stimulation (akin to the one used in the passive condition) in the baseline and block sessions, with lighter colours indicating higher firing rates. **B)** The cortical model consists of 5 units representing the different finger-selective clusters, which receive input from the periphery and are laterally connected. Thicker arrows denote stronger connections. **C)** A model fit on baseline responses based on simulated peripheral inputs cannot account for the observed results under peripheral D2 nerve block. **D)** Predicted univariate activity under D2 block for the static model (i.e. with baseline weights) against empirically observed results (see Figure 2B). Most activity is well fitted by a constant gain decrease (orange line fit to circles), but a separate mechanism is needed to account for activations arising from the blocked D2 stimulation condition (triangles). **E)** A model incorporating a global gain decrease and a selective D2 increase can reproduce the observed empirical pattern (see Figure 3). **F)** A significantly decreased Glutamate/GABA ratio following the block, as identified using MRS in the cortical hand area. The reduction in excitation/inhibition could be an empirical in vivo marker of the global gain decrease required by our model.

After fitting the model to the empirical data obtained in the baseline session (see Methods), we examined the effects of the peripheral nerve block on the cortical representations, assuming that cortical response profiles did not change. This analysis therefore tested whether the loss of peripheral input on its own might account for the observed cortical changes. As mechanical stimulation of a single finger excites tactile afferents terminating on other fingers and the palm, the loss of peripheral input from one finger could in theory affect other cortical patches. However, we found that the simulated activity pattern under anaesthesia differed from the observed one, with the static model predicting a greater collapsing of D2 representation, relative to the other fingers (square markers in Figure 4C). The observed activity patterns in the block condition therefore appear to rely on central mechanisms that mediate the effects of peripheral input loss, in addition to the distributed peripheral input.

To characterise these changes, we compared the simulated univariate activity under the static model with the empirically observed activity. This revealed that the cortical changes were driven by two simple effects. First, a global reduction in gain by approximately 25%, such that activity was reduced proportionally for all cortical units under all stimulation conditions (see fitted slope versus unity line in Figure 4D), which accounted for the majority of the block effect other than when D2 was stimulated. Second, the D2-specific maintained representation could be explained by a gain change in the D2 unit which increased its activity by approximately 25% and propagated through lateral connection to other cortical clusters (see initial model illustration in Figure 4B). Both effects together reproduced the observed cortical changes in the block condition (Figure 4E). The proposed gain changes are consistent with homeostatic mechanisms: decreased activity in the D2 unit might have elevated D2’s gain to raise this cluster’s activity, simultaneously with globally decreased gain.

The mechanisms behind the global gain changes in the model may have identifiable substrates in vivo. Enhanced excitability specific to the D2 patch may be difficult to show given the resolution of our methods. The wide-spread excitability drop for the other fingers, however, was empirically supported by changes in the neurochemical profile of S1. In a subset of participants (n=10), we successfully performed magnetic resonance spectroscopy (MRS) over the (entire) sensorimotor hand area in both baseline and block sessions. The glutamate/GABA ratio at rest, a marker of cortical excitability (Stagg et al. 2011), was reduced in the S1 hand area after block (t(9)=2.48, p=.035; Figure 4F). Although further research is warranted, this global reduction in excitation/inhibition strength provides support for a *non-specific* homeostatic plasticity mechanism that can realistically take place at the rapid timescale that applies here.

## Discussion

Here, we have shown that during a local nerve block, finger activity and selectivity was reduced globally across the entire hand area. Textbook electrophysiology findings suggest that neighbouring fingers will invade the cortical territory of a silenced finger, and this remapping in the boundaries within the hand map is taken as evidence for reorganisation. Although we were able to reproduce this remapping, we suggest that it provides a limited measure of reorganisation that should not be overinterpreted. Instead, we demonstrate that there is little evidence that voxels selective to the blocked finger at baseline increase their preference for that finger’s neighbours following deprivation, which would be expected if somatotopic remapping had taken place. Instead, we identified a significant reduction in activity and selectivity which extended well beyond the deprived finger cluster and was evenly spread across both neighbouring and non-neighbouring fingers. Furthermore, despite a drastic reduction in activity for the blocked finger in voxels that were highly selective to this finger, we found clear evidence for that finger’s persistent representation elsewhere in the hand area. Our results are consistent with recent findings in humans, showing preserved, albeit reduced, inter-finger representational structure in amputees (Wesselink et al. 2019). Furthermore, the empirical activity profiles we observed, along with the modelling results, suggest that the long-standing depiction of S1 as a set of sparsely selective topographic finger clusters paints an incomplete picture of human hand representation.

### Why was finger selectivity reduced across the hand map?

Our first finding is that differentiation between finger selectivity profiles, characterised using either local differences in net activity or multivariate representational similarity analysis, was reduced. This goes against the intuition that a local block will cause local unmasking (within cluster C2) and disinhibition within neighbouring clusters (due to reduced lateral inhibition), presumably causing increased selectivity, particularly for the neighbouring fingers. This is further supported by our computational model, suggesting that when accounting for complex inter-finger receptive fields (as will be discussed below), attenuating D2 input should not cause larger reductions to clusters C1 and C3 than to clusters selective to non-neighbouring fingers. Importantly, in the current study, the reduction in selectivity across the entire hand map was also observed during an active task, allowing us to confirm consistent input administration across sessions. To the best of our knowledge, peripheral input was comparable in the block and baseline session, save for that of the blocked finger – this will be discussed below.

The reduction in selectivity could not be accounted for solely by the reduced input from the blocked finger. Instead, as indicated by the computational model, global inhibition across the map is required to explain the observed reduction in net activity. Tonic inhibition, mediated through extra-synaptic GABA receptors, could account for this global selectivity reduction. By shifting the relative spiking threshold, receptive fields will modulate their own selectivity. Xing and Gerstein (1994) showed that rapid receptive field change following input loss can be successfully modelled using only tonic inhibition. Our observation that one finger can cause shifts to the whole hand map suggests this process is less somatotopically precise than previously assumed: the entire hand representation undergoes inhibition. This proposed mechanism fits with the decreased Glutamate/GABA ratio we have found at rest using MRS. Interestingly, we recently identified a similar whole-hand drop in selectivity, similar to what was observed here, which was observed after (able bodied) participants adapted their hand dynamics to use an additional artificial thumb [(Kieliba et al. 2020), see Ogawa et al. (2019) for similar findings in trained pianists]. Although plausible alternative theories exist for that finding (see Kieliba et al. 2020), reduced selectivity may occur more generally following atypical sensorimotor input. The mechanism behind our results may also be involved in other (rapid) representational shifts following changes in finger input, e.g. tonic inhibition following sustained tactile stimulation (Rocchi et al. 2017, Godde et al. 1996). We therefore conclude that the reduced finger selectivity observed here was aided by central plasticity mechanisms.

It is important to note that, although we have hypothesised that homeostasis is achieved through a central mechanism, plasticity mechanism occurring sub-cortically (Kambi et al. 2014) will also influence cortical representation. While we are unable to comment on the relative contribution of non-cortical mechanisms on the observed results, the altered inhibition-excitation ratio does suggest some role for S1 in this process.

### Why was the blocked finger’s representation persistent?

The observation that the blocked finger’s representation was still identifiable during blocking is also of interest. Although the peripheral signals leading to this pattern of activity mainly come from mechanoreceptors on the blocked finger (in natural contexts), some peripheral receptive fields are very large, integrating information from across most of the hand (Johansson 1978, Tommerdahl et al. 2010, even before reaching cortex: Pruszynski and Johansson 2014). These effects likely arise from the mechanical attributes of the hand, which induces a ripple of vibration following even very localised touch, which will in turn cause a specific spatiotemporal pattern of mechanoreceptor activity across the entire hand (Manfredi et al. 2012, Shao et al. 2020; see also Figure 4A). Receptive fields on the cortical level can be complex and integrate information across a variety of receptor types (Tommerdahl et al. 2010, Saal et al. 2015). As such, somatosensory input is inherently diffused across the hand. This complex cortical inter-finger representation may be also influenced by natural hand use, where inter-finger co-use is commonplace (Ingram et al. 2008) and the fingers engage in highly regular patterns of co-movement. Distributed inter-finger activity can even be induced by localised microstimulation of individual mechanoreceptive afferent units (Sanchez Panchuelo et al. 2016), suggesting this is an organisational feature of S1.

Importantly, however, the persistent response for the blocked finger in our study was higher than what the other fingers’ unblocked mechanoreceptors could have induced during the blocked finger stimulation. Our model suggests that remaining D2 information (from whichever source, see below) needs to be selectively boosted in order to produce the observed activity patterns. In other words, the global inhibition needs to be off-set with respect to residual inputs relating to D2. This could possibly be driven by a homeostatic mechanism keeping the firing rates in the deprived cortex stable. Although homeostatic plasticity is typically studied at longer timescales (Turrigiano and Nelson 2000), it may also occur at the scale of hours and less (Zenke et al. 2017), making it a potential mechanism for the changes seen here.

A more trivial explanation for the persistent D2 representation may be that we did not perform a successful nerve block. Both the diminished univariate activity of D2 in its related S1 cluster and our behavioural results showing D2 acuity at floor level indicate that input from D2 was successfully attenuated. Yet, it is likely that some D2-related residual input, though reduced, still reached S1, as clearly shown in our active condition. This is consistent with deafferentation models for studying deprivation - even when considering extreme cases of spinal cord injuries and whole-hand amputation, some rudimentary peripheral signals are likely to persist from the injured nerves (Liao et al. 2018). Indeed, in our computational model we assumed the nerve block doesn’t provide complete abolishment of all D2 inputs. While peripheral input may play a role in maintaining the representation of the amputated, deafferented or blocked body part, even then, the magnitude of the empirically observed persistent representation exceeds what would be expected from such limited input. This is supported by our model, as well as our analyses showing that the decrease in D2 dissimilarity across the hand area was not quantitatively different from the decrease observed for the other, non-blocked, fingers.

More likely, decreased peripheral drive may have been compensated by up-regulation of residual afferent, as well as non-afferent, input. First, the somatosensory cortex and the motor system are tightly linked (Kleinfeld et al. 2006, Lee et al. 2008, Adams et al. 2013) and motor commands, unaffected by our nerve block, are also routed to S1. Although this can explain why activity for the blocked finger was not reduced to zero during the active movement condition, any undetected micromovements during the passive task would not be expected to engage S1 strongly. Yet, during the active task, the global drop in selectivity was proportionally similar to that during the passive task. Second, non-afferent inputs representing cognitive (e.g., attentional) factors has been shown to modestly modulate hand-related activity in S1 (Kuehn et al. 2018, Cardini et al. 2011, Puckett et al. 2017). Pattern completion using top-down input from higher-order sensory cortex has also been shown to modulate activity in the visual cortex (Smith and Muckli 2010, Reichert et al. 2013, de Lange et al. 2018). Because the participants were informed at all times which finger was being stimulated through visual feedback, top-down processes may have been able to fill in the blocked finger’s activity. This is highly compatible with the notion of complex inter-finger receptive fields, with some residual peripheral information relating to D2 stimulation (either through some residual input, or aided by the rippling effect of mechanoreceptive responses following blocked D2 stimulation). Pattern completion could even be facilitated by the hypothesised horizontal connections between neurons within S1 (Reed et al. 2008). It is likely that all of these processes (both bottom up and top-down) contribute to the persistent representation observed here. Regardless of the specific process, the key mechanism to enable the distinct representation of D2 despite abolishment of its primary input is the distributed representation across fingers, which we believe is shaped and maintained by daily life experience (Ingram et al. 2008, Dempsey-Jones et al. 2016).

### Was Merzenich wrong?

While we find wide-spread changes in finger selectivity for hand representation following localised deprivation, we do not observe any changes in the overall inter-finger representational pattern. That is, we find a shrinkage of the representational structure rather than the kind of reorganisation typically attributed to local input loss in S1 (see Introduction). This is consistent with our findings in amputees, but inconsistent with previous electrophysiological studies of animals. It is important to consider the methodological differences between our present study and these classical studies, which could have contributed to the diverging results. Contrary to the time scale of these classical studies (hours to days), the speed at which our effect occurs (within 1-2 hours) indicates that it is unlikely for the observed changes to be driven by substantial anatomical or synaptic plasticity (see also Rasmusson and Nance 1986). Still, enlargement of receptive fields following deprivation has been replicated for amputation (Turnbull and Rasmusson 1990, Calford and Tweedale 1991, Kolarik et al. 1994) and anaesthesia (Calford and Tweedale 1991, Panetsos et al. 1995, Faggin et al. 1997), sometimes after less than one hour post-intervention. While each neuron’s suprathreshold responses are highly plastic, the widespread arborisation of afferent connections providing the excitatory input, is more stable (Landry and Deschenes 1981, Garraghty et al. 1994, Rausell and Jones 1995, Calford 2002). Inhibitory networks dynamically determine which responses cross the threshold, creating the traditional receptive field. It is possible we did not see expanded receptive fields in our study, i.e. no increase in activity for neighbouring fingers in the D2 cluster, because fMRI pools many cells’ response profiles, so our observed distributed activity, even at baseline, might have already been less than fully inhibited (and thus measurable).

In our view, the many examples in the literature for the behavioural state and input statistics presumably affecting how inhibition shapes receptive fields, do not necessarily indicate reorganisation, i.e. a change away from a norm. Instead, they may show that topographical representation in S1 dynamically reflects task demands (Weinberger 1995, Calford 2002), rather than merely our rigid body. For example, while enhanced tactile stimulation of fingers can cause receptive field changes, this process is both rapid and reversible (Godde et al. 1996), suggesting that such changes are within the hand area’s representational dynamic range. The importance of input history for the response profile of cortical representations was also stressed by Merzenich et al. (1987). While it is possible that, at a longer timescale, other mechanisms in line with Hebbian plasticity could produce more substantial adult reorganisation (Dykes 1997, Feldman 2009), we and others find that, beyond the dynamic representational features described above, adult primary sensory cortex shows remarkable stability (Smirnakis et al. 2005, Makin and Bensmaia 2017).

## Conclusion

Here, we have shown that despite drastic and highly localised changes to the periphery, cortical hand representation shows relatively homogeneous consequences. Our results indicate that the somatosensory cortex is not a patchwork of isolated clusters representing individual digits. Complex interactions exist between neighbouring representations in the periphery, cortex and in-between, all of which ought to be taken into consideration before invoking plasticity mechanisms to explain observed remapping. If this interconnectedness comes with the ability to complete missing input in certain contexts, as our findings suggest, new opportunities are opened up for restorative applications and BCI control.

## Methods

### Participants

Fifteen healthy volunteers (6 female, age = 26.44 ± 1.04) participated in this study. In addition, one volunteer (P16) participated in session 1 but did not complete session 2; their imaging data has not been included but we did include the collected behavioural data. All participants but one were right handed and all experimental tasks were performed using the right hand. All participants gave written informed consent and ethical approval for the study was obtained from the Health Research Authority UK (13/SC/0502).

One participant was excluded from the passive multivariate task analysis since they were identified as an outlier (score ≫3 standard deviations below the group mean in the baseline session). This participant was included in the active condition where their scores were considered to be within normal group variance (0.3 standard deviations above group mean). In addition, due to multiple technical challenges with data acquisition (due to the complex MRI protocols, the induction of the nerve block, the time-consuming behavioural testing) and due to the exceedingly long testing session, we sustained some impartial datasets. Fourteen participants took part in the post-scan grating orientation task, but two missed one of the two sessions. Thirteen participants had resting-state data for both sessions, and ten participants had stable magnetic resonance spectra in both sessions (see below for more details).

### Experimental procedures

Participants attended two sessions that were similarly structured (see Figure 1A). A finger nerve block was administered in one of the two sessions (in counter-balanced order between participants). In both scan sessions participants completed an active and a passive task, a resting-state, and spectroscopy scans. In the baseline session, participants also performed a functional localiser task which was used to define the finger-selective clusters.

### Procedures outside the scanner

#### Pharmacological nerve block

Seven participants received the nerve block in the first session and eight received it in the second session. The nerve block consisted of a mixture of 2 mL of lidocaine 2% and 2 mL of Bupivacaine hydrochloride 0.5%, allowing for both a rapid onset of the anaesthetic effect and a stable effect of 5-8 hours (depending on blood circulation). The solution was injected by a trained medical professional around the base of the right index finger and each side of the finger was injected with approximately half of the solution. This thereby formed a ring block, anaesthetising the entire finger.

#### Psychophysical testing

We used a range of tactile acuity and sensitivity assessments to verify the perceptual effectiveness of the nerve block. All measures were carried out during both the baseline and block sessions. To test acuity, an experimenter applied a plastic domed grating with a grating width of 3.5 mm to the glabrous surface of the right distal index finger perpendicularly for a period of approximately 1s. The grating was oriented vertically or horizontally, randomly over 20 trials, and participants reported the perceived orientation via a mouse click (with their left hand). This provided a relatively quick means to probe tactile acuity and was carried out at four time points: an initial test shortly after the injection, immediately prior to the scan, immediately after the scan and after the more precise post-scan sensitivity tests (see Figure 1A). Performance above 70% correct in the pre-scanning test of the block session was an a priori criterion for exclusion. Based on this criterion, all participants were successfully blocked. We also tested the adjacent D3 to confirm that the effects of the nerve block were localised.

After the injection (but prior to the scan), we also assessed tactile sensitivity using Von Frey hairs. Two hairs of different forces (2.0 mN and 8.0 mN) were applied to the glabrous surface of the distal index finger. Participants were asked to indicate which force felt stronger in each trial, over 20 trials. On each trial, the filament was pressed perpendicularly against the fingertip for approximately 2s.

After the scan, we estimated tactile acuity thresholds more precisely for the index, middle and ring fingers. Plastic domed gratings with five different grating widths (0.5, 1.0, 1.5, 2.5, 3.5 mm) were briefly applied to each of the finger’s distal pad in either horizontal or vertical orientation, over 20 trials per grating width and finger. Participants were asked to indicate the orientation of the grating with a mouse click of the left hand. Participants were blindfolded throughout the procedure. The order in which the gratings were presented was randomised. The trials were grouped in two blocks interspersed by a short break. To encourage engagement, participants received intermittent auditory feedback on their performance (percentage correct) over headphones.

### MRI tasks

#### General procedures

In a single session, participants completed an active and a passive task in the scanner (in counterbalanced order between participants), as well as an anatomical scan, a resting state scan and a MR spectroscopy (MRS) scan (also performed during rest). The active and passive tasks were completed over four separate runs (for each task), as described in Sanders et al. (2019).

The scan protocol was identical for the passive and active task. Each run comprised of individual finger blocks for each of the five fingers (one tap per second, 12s blocks) of the right hand, as well as no movement (rest) blocks lasting 12 or 24s. Each condition was repeated 3 times in a semi-counterbalanced order within each run, here termed ‘random design’. Each run totalled 118 volumes and comprised a different block order.

We also conducted a functional localiser before the active task in the baseline session to independently identify finger-selective regions of interest (here termed clusters C1-C5). This localiser was also organised into finger-specific blocks, but with a set inter-finger sequence design (‘travelling wave design’; Kikkert et al. 2016, Kolasinski et al. 2016, Wandell et al. 2007, Sanchez-Panchuelo et al. 2010, Mancini et al. 2012, Zeharia et al. 2015). Each block lasted 8 seconds. Two runs were acquired, with a reversed order from each other, each consisting of 108 volumes, covering 5 cycles around the hand.

#### Passive task

In the passive stimulation task, we asked the participants to rest their right hand in a comfortable, supine position on a foam cushion. An experimenter stimulated each finger by tapping a plastic probe against the distal pad of the finger. Such manual stimulation is commonly used in somatosensory studies to limit “contamination” from the motor system and has previously shown to robustly activate finger maps (Martuzzi et al. 2014). The experimenter was instructed through headphones. Any slight variations produced by the experimenter were meant to account for variations in the active task (below), and were not likely to differ between the baseline and block sessions. During the passive task, participants were shown dots flashing synchronously with the tactile stimulation to indicate when and where a touch occurred. To promote engagement across the duration of the task, double taps were administered sporadically (one double-tap per finger condition in each session). Participants were asked to press a button with their left hand when they felt a double tap. Participants correctly identified these catch trials in 93.3% of the cases in the baseline session (excluding D2 trials). This percentage was 94.2% in the block session. There was no significant difference in double tap detection between (non-blocked) fingers (F(3,112)=1.97, p=.123) or between sessions (F(1,112)=0.10, p=.748). Detection of D2 double taps was impaired in the block session (66.7%) compared to baseline (98.3%, t(14)=3.68, p=.002).

#### Active task

The active task was a visually cued (motor) task. In an intact sensorimotor system, movement recruits a combination of peripheral receptors, encoding a range of somatosensory modalities (e.g. surface and deeper mechanoreceptors; proprioceptors), as well as efferent information from the motor system. Using an active task, we have previously shown high consistency of S1 finger topography across multiple scanning sessions (Kolasinski et al. 2016, see also Ejaz et al. (2015) for validation using RSA). Participants were presented with five vertical bars, corresponding to the five fingers, shown on a visual display projected into the scanner bore. To cue the participant which finger should be moved, the bar corresponding to this finger changed (i.e. by flashing in a different colour).

The participants performed the tasks well. The instructed finger produced the strongest press force in 94.6% of the trials (92.2% in the worst participant). Consequently, there was a clear difference in average force output between the instructed and non-instructed fingers: In the baseline session, 1.44 N (+/− 0.09 SEM) for the instructed finger and 0.27 N (+/− 0.03) for non-instructed fingers; and in the block session, 1.39 N (+/− 0.07) and 0.24 N (+/− 0.03) respectively. There was no difference in force output between sessions (F(1,140)=0.28, p=.596). This was also the case when only D2 output was compared (t(14)=1.31, p=.211), suggesting any differences between sessions are not due to impaired motor performance.

#### Finger-selective cluster localiser

The travelling wave protocol involves a set finger cycle. This approach is particularly useful for identifying voxels that show an enhanced response to one finger compared to all other fingers, and has previously been used to identify S1 finger somatotopy (e.g. Kolasinski et al. 2016). Participants used the same keyboard and visual display as in the active task. Two separate runs, with a reverse orders of conditions (i.e. a forward and a backward cycle), were used to overcome potential order-related biases due to the sluggish haemodynamic response. In the forward run, the order of finger blocks cycled from finger 1 to finger 5 (D1-D2-D3-D4-D5) whereas a reverse order of finger blocks was used in the backward run (D5-D4-D3-D2-D1). In each run, the cycle was repeated five times with no rest periods in between. During the cycles, each finger was moved 8 times (at 1Hz) before the instructions for the next finger were shown. As in the active task, the finger to be used in the upcoming block was visually cued at the start of each block, followed by 8 finger presses of that same finger.

#### Resting state scan

Participants were instructed to keep their eyes open and gaze at a fixation cross. Otherwise, they were instructed to let their mind wander and not think of anything in particular. 150 volumes were acquired.

### Tactile perceptual analysis

For the sensitivity checks of grating orientation judgement (Fig 1B-C), the four measurements were grouped into pre-scan (2 tests) and post-scan (2 tests) measurements. Session comparison was done on the average percentage of correct responses.

For the detection performance with the Von Frey hairs (Fig 1D), raw accuracy data was Z-normalised and d prime (d’) scores were calculated (hits minus false alarms).

Tactile psychophysical thresholds were determined separately for each finger/session, using standard procedures (e.g. Dempsey-Jones et al. 2019). In short, this was done by plotting accuracy as a function of grating size across all levels of stimulus difficulty and fitting a Weibull curve using a least-squares function in MATLAB (two free parameters; gamma = 0.05, lambda = 0). The threshold for this psychometric function was interpolated from the grating size estimated to yield 82% accuracy. We also extracted the slope and goodness of fit values (R2) for each curve fit (slope is taken as the steepness of the psychometric function at the threshold point, and the R2 represents how well the psychometric function represented the data, i.e. how close the data points are to the line). Overall, the psychometric functions predicted the data with relative accuracy (average R2 = .76, SEM = .02). Thresholds were compared across fingers and sessions.

### MRI acquisition & pre-processing

#### MRI acquisition

All MRI measurements were acquired using a Siemens 7 Tesla Magnetom scanner with a 32-channel head coil. Task fMRI data was acquired using a multiband EPI sequence with an acceleration factor of 2 (Moeller et al. 2010, Ugurbil et al. 2013). A limited field-of-view was used for fMRI acquisition, consisting of 56 slices of 1mm thick, centred over S1 with a 192×192mm in-plane FOV (TR 2000ms, TE 25ms, FA 85deg, GRAPPA 3). This resulted in a spatial resolution of 1mm isotropic. A whole brain anatomical T1-weighted image was also collected with 1mm isotropic spatial resolution (TR 2200ms, TE 2.82ms, FA 7deg, TI 1050ms).

1H magnetic resonance spectroscopy (MRS) data was acquired and pre-processed as described in (Lunghi et al. 2015). A 2 × 1 × 1 cm voxel was placed manually over the hand knob in S1 using the collected T1-weighted anatomical scan. Three guidelines were followed to motivate correct placement (in order of importance): 1) the voxel avoided the dura mater, to prevent signal issues; 2) the voxel was placed posterior to the central sulcus, to limit the influence of M1; and 3) the voxel was placed as superior as possible to focus on the hand region. Due to data acquisition errors, data from three participants has been excluded from analysis. Two further participants were excluded from the analysis due to unreliable GABA quantification in one of the sessions (Cramér-Rao lower bounds higher than 50%). As such, the placement and resulting data quality was sufficient in both sessions to produce reliable spectra for 10 participants.

#### MRI pre-processing

All MRI data pre-processing and analysis was carried out using FMRIB Software Library (FSL; version 6.0) as well as Matlab scripts (version R2016a) which were developed in-house. Surface reconstruction was carried out using Freesurfer [(Dale et al. 1999); www.freesurfer.net] and results from the task and travelling wave analysis were projected onto the cortical surface for visualisation purposes using Connectome Workbench software (www.humanconnectome.org).

Standard pre-processing steps were carried out for the fMRI data using FSL (Jenkinson et al. 2012). FSL’s Expert Analysis Tool (FEAT) was used to carry out motion correction (using MCFLIRT; Jenkinson et al. 2002), brain extraction (BET; Smith 2002), spatial smoothing of all fMRI data using a 1mm full width at half maximum (FWHM) Gaussian kernel and high pass filtering using a cut-off of 100s. The output from the MCFLIRT analysis was visually inspected for excessive motion (defined as >1mm absolute mean displacement). No participants had an absolute mean displacement greater than 1mm.

#### Image registration

For each participant, a mid-space was calculated between the four active and four passive runs, i.e. the average space in which the images are minimally reoriented. Each scan was then aligned to this session mid-space using FMRIB’s Linear Image Registration Tool (FLIRT; 6 DOF; Jenkinson and Smith 2001, Jenkinson et al. 2002). The two runs of the functional localiser were also registered to the mid-space of the baseline session (but first to each other). The structural scans from both sessions were also combined by finding a mid-space. The functional mid-spaces from both sessions were registered to this anatomical mid-space using FLIRT together with manual adjustments to ensure an accurate co-registration of the central sulcus (specifically, the “hand knob”). Once co-registration was satisfactory, all functional scans across both sessions were aligned to this anatomical mid-space.

### Task-based fMRI analysis

#### Active and passive tasks

A voxel-based general linear model (GLM) analysis was carried out on each of the 16 runs (four active and four passive per session) using FEAT, to identify activity patterns for each finger condition. The design was convolved with the double-gamma haemodynamic response function, as well as its temporal derivative. For each run, eleven contrasts were set up: each finger versus rest, each finger versus all other fingers and all fingers versus rest. For univariate analysis, the estimates from the four active/passive runs were then averaged voxel-wise for each participant using a fixed effects model, creating eleven main activity patterns for each task. These contrasts were used for the follow-up analysis described below.

#### Localiser & Cluster definition

The travelling wave task described above was used in order to identify finger-specific voxel clusters. The analysis was carried out as in Kikkert et al. (2016). In short, a reference model was created using a convolved hemodynamic response function to account for the hemodynamic response. This model consisted of an 8 second ‘on’ period followed by a 32 second ‘off’ period to model the movement block of one finger for one cycle. The model was shifted 20 times (i.e. the amount of lags/TRs of a single movement cycle) in order to model one entire movement cycle (which lasted 40s), thus resulting in 20 reference models. This was repeated five times to model the five cycles in each run. Each of these reference models was then correlated with each voxel’s pre-processed BOLD signal timecourse. This resulted in cross correlation r-values for each voxel, which were standardized using the Fisher’s r-to-z transformation. Lags were assigned to each finger (four lags per finger) in order to average the r-values across runs for each voxel. This resulted in a r-value for each finger which was further averaged across the forward and backward runs. To define finger specificity, each voxel was assigned to one finger using a ‘winner-takes-all’ approach. This was done by finding the maximum for each voxel across the five averaged r-values, and assigning the voxel to the corresponding finger.

To correct for multiple comparisons, a false discovery rate (FDR; Benjamini and Hochberg 1995) threshold (q < 0.01) was applied to each finger individually (as in Kikkert et al. 2016). The resulting FDR corrected finger-specific voxels were then used to create finger-specific clusters showing strong finger selectivity for each of the fingers. This was done by using an anatomically defined mask of the S1 hand area which was defined for each participant based on a Freesurfer structural segmentation of S1 subdivisions. Brodmann Areas 3a, 3b and 1, spanning a 2cm strip dorsal/ventral to the anatomical hand knob were included in the mask (Wesselink et al. 2019). The finger specific activity under this mask was used to create finger-specific S1 clusters (C1-5, see e.g. Figure 2). This approach allowed us to identify finger-specific S1 clusters on an individual participant basis, which were used for further analysis of the active and passive tasks. Clusters are not necessarily contiguous. The size of these clusters varied considerably between participants, as reported in. Table S1. Finally, the entire S1 hand mask was also used as its own region-of-interest for the winner-takes-all (remapping) analysis reported in Figure 2B-C and the multivariate analysis reported in Figure 3B-C (excluding highly selective voxels in the clusters).

#### Univariate selectivity analyses & winner-takes-all maps for the main tasks

To examine the distribution and selectivity of finger activity in the active/passive tasks, average activity was calculated for each finger in each finger cluster. Activity was calculated separately for each finger, condition, and session. As a measure of selectivity, we subtracted in each cluster the average activity of the non-target fingers from that of the target finger (i.e. the finger that the cluster is selective for). For this analysis, as reported in the main text, activity of D2 and activity in cluster C2 were ignored, so each cluster had 3 non-target fingers.

In order to create winner-take-all maps for the passive task, we first projected the five finger-versus-rest contrasts (averaged across runs) from the passive task onto the cortical surface. Within the S1 hand area (see above), each voxel was either assigned to the finger that evoked the strongest BOLD response, or assigned as non-active if the maximum response did not exceed 0. In order to simulate classical analysis carried out following amputation, this analysis was repeated while ignoring D2, i.e. voxels could only be assigned to one of four fingers. To examine whether rapid invasion occurred in the deprived cortical territory, we compared the number of voxels assigned to each finger within D2-selective cluster C2.

#### Representational similarity analysis

We used representational similarity analysis (Kriegeskorte et al. 2008) to assess the multivariate relationships between the activity patterns generated across fingers and tasks. The (dis)similarity between activity patterns within the S1 hand mask was measured for each finger pair using the cross-validated Mahalanobis distance (Walther et al. 2016). The activity patterns were pre-whitened using the residuals from the GLM and then cross-nobis distances were calculated for each task (active/passive) separately, using each pair of imaging runs and averaging those results. Due to cross-validations, the distance value is expected to be zero (but can go below) if two patterns are not statistically different from each other. Otherwise, greater distances indicate larger differences in multivariate representation. The analysis above produced 10 distance values (one for each finger pair) per task/region of interest, forming a representational dissimilarity matrix (RDM) per participant. These values can statistically be compared to 0 (i.e. representing no difference between conditions or rest; see *Statistics* for more details). This analysis was repeated for three regions of interest: within each finger-selective cluster (Figure 3A), across the five finger-selective clusters (Figure 3B) and across the entire hand area, after excluding the finger-selective clusters (Figure 3C).

As an aid to visualise the RDMs (and not used in any statistical analysis), we also performed multidimensional scaling (MDS). This analysis projects the higher-dimensional RDM into a lower-dimensional space while preserving the inter-finger distances as accurately as possible (Borg and Groenen 2005). Rest was included as an extra condition (with uniform activity of 0), so finger-specific differences could be separated from differences from rest. MDS was performed on individual data and averaged across participants after Procrustes alignment (without scaling) to remove any arbitrary rotations introduced by MDS. The two dimensions along which the data was projected showed the maximal between-finger variance, reflecting differences between fingers rather than non-finger-specific variation from rest.

### Magnetic resonance spectroscopy

1H magnetic resonance spectroscopy (MRS) was acquired using 2×1×1 voxel placed manually over the hand knob in S1, using the collected T1-weighted anatomical scan. Three guidelines were followed to motivate correct placement (in order of importance): 1) the voxel avoided the dura matter, to prevent signal issues; 2) the voxel was placed posterior to the central sulcus, to limit the influence of M1; and 3) the voxel was placed as far superior as possible, to focus on the hand region.

Spectra were measured with a semi-adiabatic localization by adiabatic selective refocusing (semi-LASER) sequence (TE=36ms, TR = 5s, 64 averages) with variable power RF pulses with optimized relaxation delays (VAPOR), water suppression and outer volume saturation (Deelchand et al. 2015, Oz and Tkac 2011). Unsuppressed water spectra acquired from the same volume of interest were used to remove residual eddy current effects and to reconstruct the phased array spectra (Natt et al. 2005).

MRS metabolites were quantified using LCmodel. The model spectra of alanine (Ala), aspartate (Asp), ascorbate/vitamin C (Asc), glycerophosphocholine (GPC), phosphocholine (PC), creatine (Cr), phosphocreatine (PCr), GABA, glucose, glutamine (Gln), glutamate (Glu), glutathione, lactate (Lac), *myo*-Inositol (*myo*-Ins), NAA, N-acetylaspartylglutamate, phosphoethanolamine (PE), scyllo-Inositol (*scyllo*-Ins) and taurine were generated based on previously reported chemical shifts and coupling constants by the GAMMA/PyGAMMA simulation library of VeSPA (Versatile Simulation, Pulses and Analysis) according to a density matrix formalism. Simulations were performed with the same RF pulses and sequence timings as those on the 7T system in use. Resonances were assigned according to their known 1H chemical shift along the spectrum (x-axis, in parts per million). The T2 relaxation of tissue water content (80 ms; Deelchand et al. 2015) was taken into account in the LCmodel fitting. Absolute neurochemical concentrations of GABA and Glutamate were extracted from the spectra of the greater S1 hand area while correcting for voxel tissue content (Lunghi et al. 2015). Metabolites quantified with Cramér-Rao lower bounds higher than 50% (estimated error of the metabolite quantification) were classified as not detected. The Glutamate/GABA ratios for all participants were compared across sessions using a paired t-test.

### Resting state scan

Following the same pre-processing pipeline as for the task fMRI (see above), the mean time course of the resting state fMRI scan was extracted from the S1 region of interest. The standard deviation of the timeseries over time was compared between sessions. Less signal variation would suggest diminished excitability that is not task-specific.

### Modelling details

Peripheral inputs were generated using TouchSim (Saal et al. 2017), a model that reconstructs typical afferent brain activation to stimuli placed across the hand. For the baseline condition, modelled tapping on each of the five fingertips generated input mimicking that observed in the passive fMRI task. The nerve block was simulated in the model by reducing the activity of all afferents in D2, for all stimulus profiles, to 20% of baseline inputs. Thus, we assume some small residual activation. Peripheral input to a cortical cluster was pooled for each finger.

The cortical model consists of a feedforward part, where each cortical cluster receives input from all fingers. It further involves a set of lateral connections, where each cortical cluster excites or inhibits other clusters. The strength of the lateral excitation and inhibition is determined by the activity of the cortical cluster. Thus, at each timestep *t*, the activity of a single cortical cluster can be calculated as:

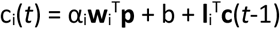

where *c*_*i*_(*t*) is the activity of cortical cluster *i* responses at timestep *t*, α is a gain factor, **w** is a weight vector specifying each finger’s connection strength to the cortical cluster, **p** is the vector of peripheral inputs from each finger, b is a scalar offset, **l** is the lateral connection field and **c**(t-1) is the vector of cortical activity from the previous time step. To simulate cortical activity under peripheral stimulation, we iterated the above equation until the cortical cluster activity had settled, usually within 6 time steps.

For the baseline model, all gain factors α were set to 1. The lateral connection **l**_ij_ between clusters *i* and *j* depended on the distance between the cortical clusters alone and therefore each cortical cluster had the same lateral connection pattern, only shifted spatially. In the model presented here, we assumed short-range excitation and long-range inhibition. Such a lateral connection pattern is frequently assumed in cortical modelling and can indeed recreate the canonical RSA hand representation commonly observed in S1. However, we note that shifting the “rest state” against which cortical activity is expressed, will also change the lateral connectivity pattern and, for example, lead to inhibition between neighbouring clusters. We cannot disambiguate between models that only differ by what “rest state” is assumed, but all such models require both a global and a D2-specific gain shift to mimic the empirical results under block, so our main results are unaffected. In Figure 4C-E, we increased the baseline to allow for the typical pattern of lateral connectivity shown in Figure 4B. The model parameters **w**_i_ and b were fitted to the empirical data in the baseline session, using the TouchSim generated finger responses. Feedforward weights between the fingers and cortical units were initially fit using multiple regression. Then, an iterative process was used to adjust the weights based on the settled activity after multiple lateral updates. RDMs were calculated using the Euclidean distances between the responses of the modelled cortical units for each finger stimulation.

Using the feedforward weights and lateral connectivity pattern generated using baseline session input, we then calculated the result of D2 block without any additional cortical changes (i.e. the static model). As this led to a poor fit, we let the gain parameters α vary to test whether homeostatic mechanisms might account for the observed changes. We found that two changes were required in order to reproduce the experimental findings. First, a global reduction in the activity of all fingers (α = 0.75), and secondly, an increase in the D2 cortical cluster gain (α_2_ = 1.25). This increase enables widespread changes in the activity of all clusters via propagation through the lateral connections and stabilizes the cortical representation of D2.

#### Statistics

Unless stated otherwise, statistical comparisons were done using (paired or one-sample) two-tailed t-tests and (RM-)ANOVA. To estimate tactile acuity and sensitivity thresholds, we used GEEs (general estimating equations). To confirm the null-hypothesis for key non-significant results, Bayesian statistics was carried out, as implemented in JASP (JASP Team 2020; using as a prior as Cauchy distribution centred on 0 with a width of .707). BF10 under .33 was considered as substantial evidence in favour of the null (Wetzels et al. 2011, Dienes 2014). The significance of rank correlations between the canonical representational finger pattern and non-target fingers in each cluster was calculated by comparing the sample mean against that of permuted labels.

## Supplementary Section

### Univariate finger selectivity including D2

When D2 was included among non-target fingers, there was a significant decrease in selectivity across the finger clusters in the active task (F_(1,112)_=5.00, p=.0273), but not in the passive task (F(1,112)=3.62, p=.059). The session x cluster interaction was not significant in either task (F_(3,112)_=0.61, p=.607; and F(3,112)=0.41, p=.745, respectively).

### Supplementary figures and tables

**Figure S1:**
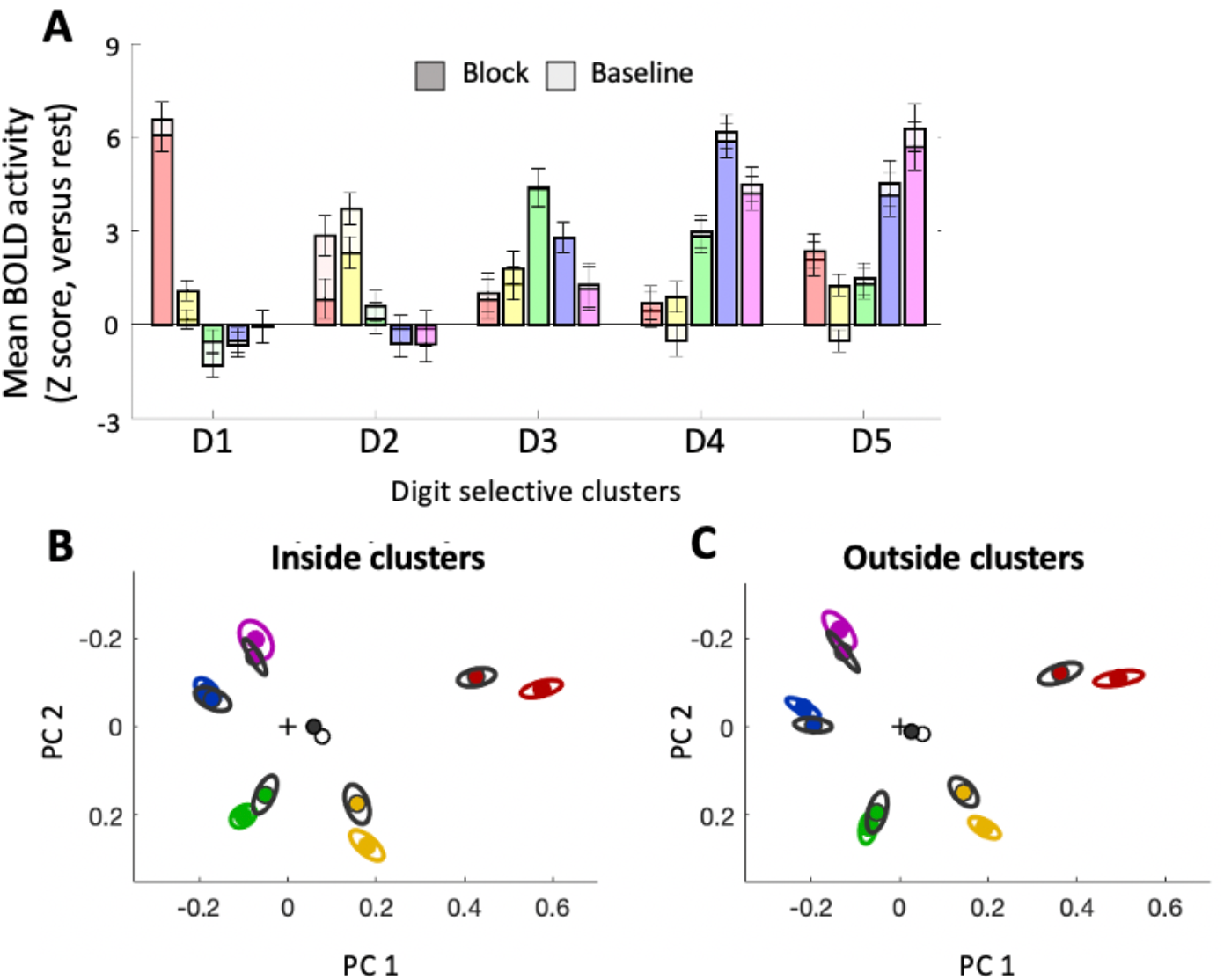
Univariate and multivariate results from the active condition. Related to Figures 2B and 3B-C. All other figure annotation is as detailed in the main figures. A) In the block session, active stimulation elicited positive activity in the deprived cluster C2 (μ=2.13; t_(14)_=4.40, p=.001), but activity was significantly reduced compared to the baseline session (t_(14)_=−2.22, p=.044). Mean activity in cluster C2 for the neighbouring fingers D1 and D3 was decreased in the block session (t_(14)_=−2.51, p=.025). B-C) In line with the passive results, dissimilarity from rest during the active task shows a strong effect of session (B: F_(1,140)_=10.7, p=.001; C: F_(1,140)_=15.92, p<.001) and no session x finger interaction (B: F_(4,140)_=0.21, p =.930; C: F_(4,140)_=0,28, p=.891).

**Figure S2:**
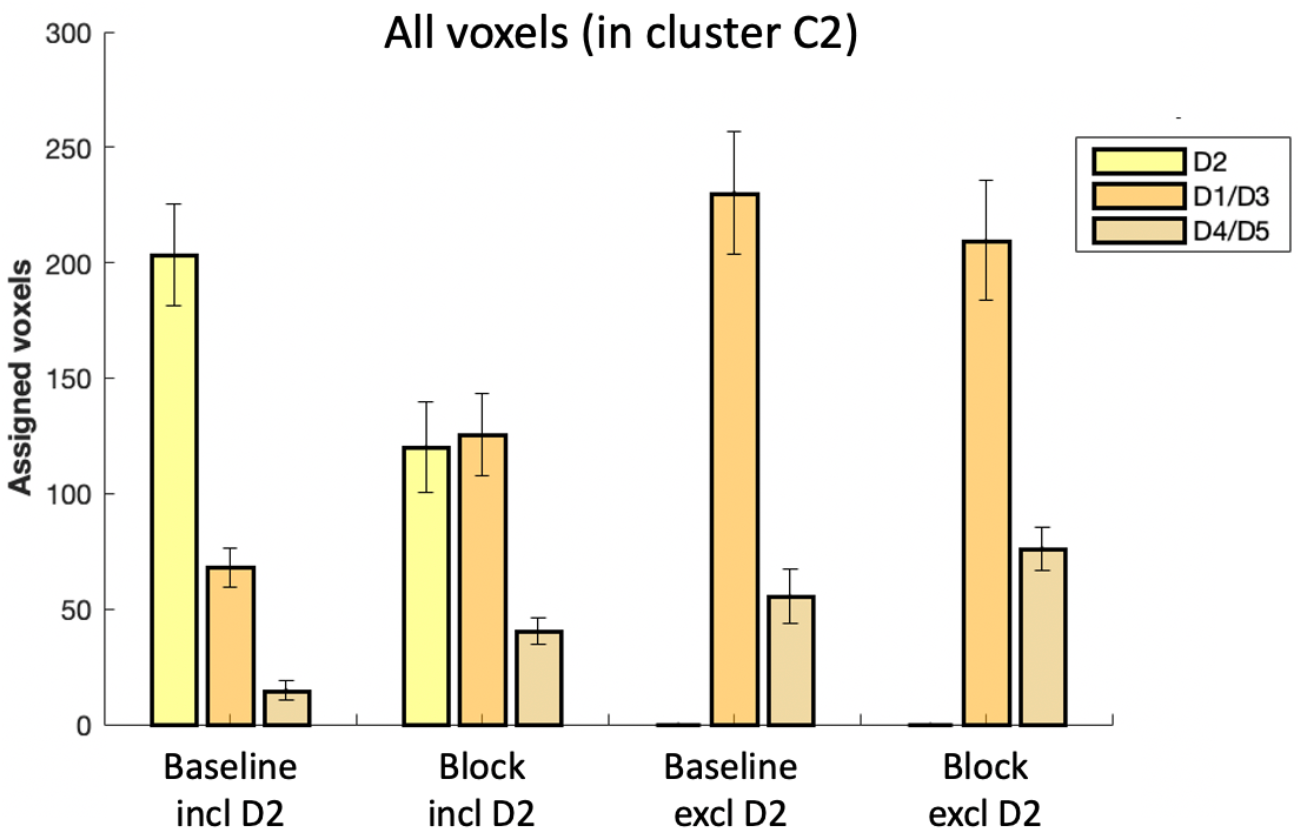
Quantification of ‘remapping’, using unthresholded voxels. Related to Figure 2D. All other figure annotations are as detailed in the main figure. As with the thresholded voxels, when D2 was ignored in both sessions (i.e. excluded from the winner-takes-all analysis), no significant difference in neighbouring fingers remapping was found between the baseline and block sessions (t(14)=1.61, p=0.131).

**Figure S3:**
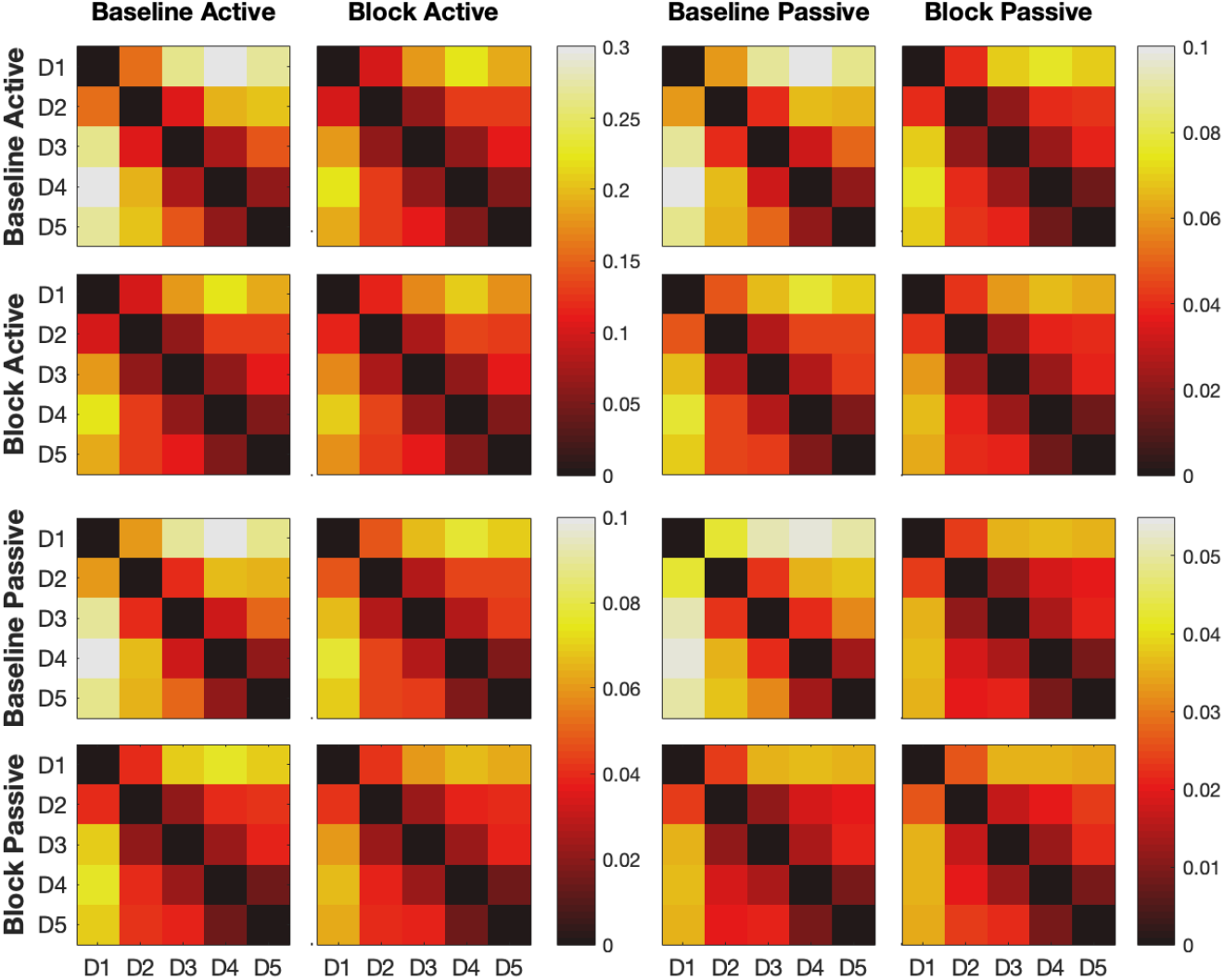
Representational dissimilarity matrix for all conditions, entire hand map. Related to Figure 3. Colours indicate Mahalanobis distance (arbitrary unit), please note scales vary between blocks. Each block compares finger activity patterns across conditions and sessions, assuming equal “rest state” (i.e. no activity) between sessions.

**Table S1.** Number of voxels per cluster. For the RSA analysis of individual clusters (e.g. in Figure 3A), clusters with fewer than 50 voxels were excluded; these have been highlighted in red.

|  | C1 | C2 | C3 | C4 | C5 | Total hand area |
| --- | --- | --- | --- | --- | --- | --- |
| P1 | 481 | 155 | 43 | 324 | 211 | 12818 |
| P2 | 456 | 229 | 268 | 391 | 197 | 7839 |
| P3 | 468 | 151 | 388 | 380 | 124 | 7080 |
| P4 | 284 | 290 | 252 | 358 | 54 | 6054 |
| P5 | 610 | 213 | 219 | 598 | 241 | 5483 |
| P6 | 694 | 425 | 221 | 42 | 368 | 6461 |
| P7 | 136 | 336 | 276 | 260 | 66 | 5903 |
| P8 | 351 | 96 | 162 | 225 | 1 | 5778 |
| P9 | 92 | 176 | 50 | 154 | 253 | 6150 |
| P10 | 719 | 270 | 65 | 292 | 302 | 8411 |
| P11 | 374 | 398 | 304 | 609 | 235 | 7100 |
| P12 | 194 | 187 | 109 | 284 | 212 | 5350 |
| P13 | 230 | 76 | 21 | 331 | 249 | 5802 |
| P14 | 368 | 140 | 54 | 416 | 127 | 5629 |
| P15 | 771 | 235 | 222 | 172 | 83 | 5271 |

**Table S2.**
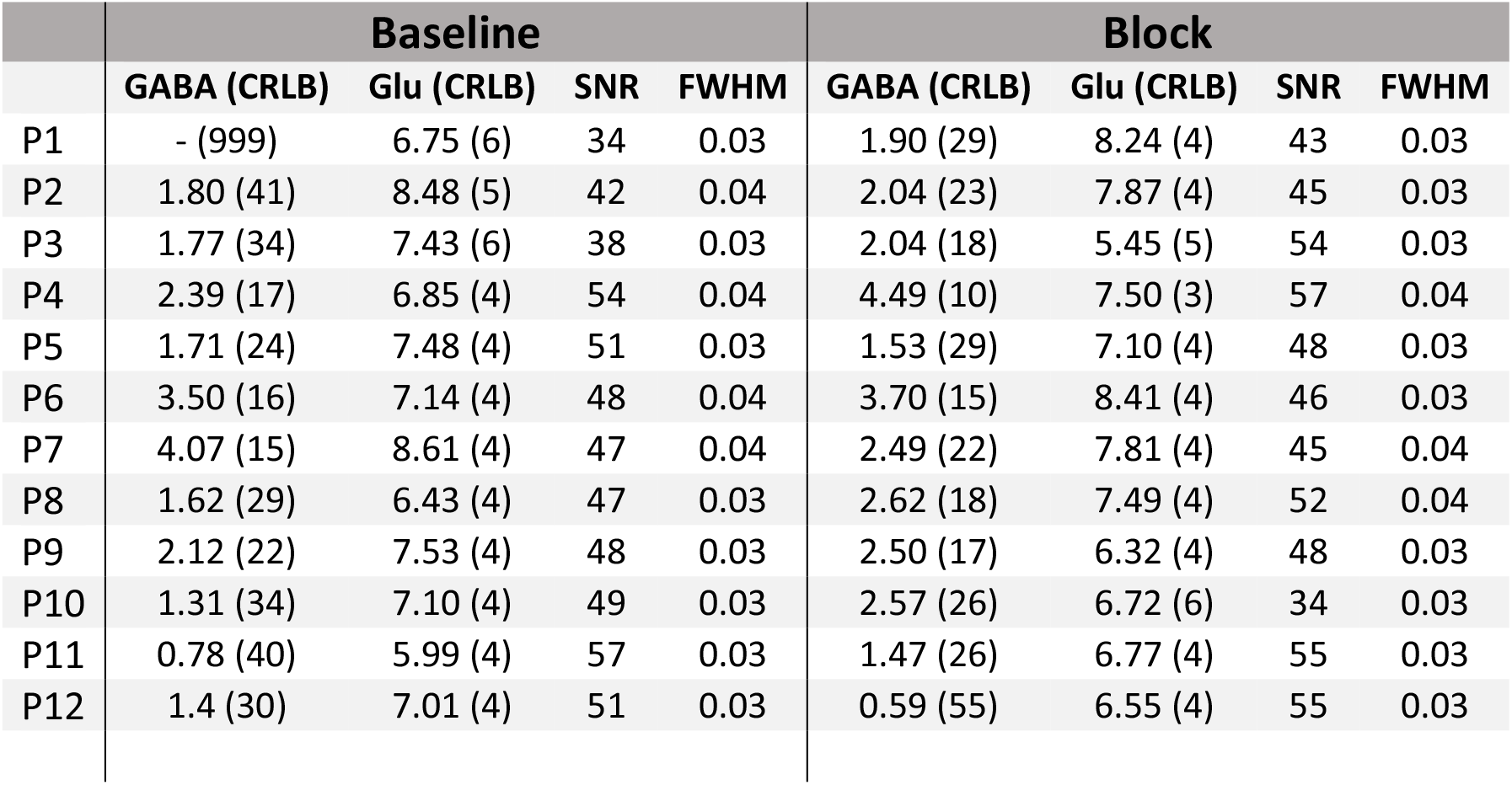
GABA and Glutamate (GLu) values. GABA and Glu estimates of all individual subjects, together with their Cramér-Rao lower bounds (CRLBs), signal to noise ratio (SNR) and full width half maximum (FWHM). Note that subjects P1 and P12 were excluded from the analysis, due to unreliable GABA readings.

|  | Baseline |  |  |  | Block |  |  |  |
| --- | --- | --- | --- | --- | --- | --- | --- | --- |
|  | GABA (CRLB) | Glu (CRLB) | SNR | FWHM | GABA (CRLB) | Glu (CRLB) | SNR | FWHM |
| P1 | - (999) | 6.75 (6) | 34 | 0.03 | 1.90 (29) | 8.24 (4) | 43 | 0.03 |
| P2 | 1.80 (41) | 8.48 (5) | 42 | 0.04 | 2.04 (23) | 7.87 (4) | 45 | 0.03 |
| P3 | 1.77 (34) | 7.43 (6) | 38 | 0.03 | 2.04 (18) | 5.45 (5) | 54 | 0.03 |
| P4 | 2.39 (17) | 6.85 (4) | 54 | 0.04 | 4.49 (10) | 7.50 (3) | 57 | 0.04 |
| P5 | 1.71 (24) | 7.48 (4) | 51 | 0.03 | 1.53 (29) | 7.10 (4) | 48 | 0.03 |
| P6 | 3.50 (16) | 7.14 (4) | 48 | 0.04 | 3.70 (15) | 8.41 (4) | 46 | 0.03 |
| P7 | 4.07 (15) | 8.61 (4) | 47 | 0.04 | 2.49 (22) | 7.81 (4) | 45 | 0.04 |
| P8 | 1.62 (29) | 6.43 (4) | 47 | 0.03 | 2.62 (18) | 7.49 (4) | 52 | 0.04 |
| P9 | 2.12 (22) | 7.53 (4) | 48 | 0.03 | 2.50 (17) | 6.32 (4) | 48 | 0.03 |
| P10 | 1.31 (34) | 7.10 (4) | 49 | 0.03 | 2.57 (26) | 6.72 (6) | 34 | 0.03 |
| P11 | 0.78 (40) | 5.99 (4) | 57 | 0.03 | 1.47 (26) | 6.77 (4) | 55 | 0.03 |
| P12 | 1.4 (30) | 7.01 (4) | 51 | 0.03 | 0.59 (55) | 6.55 (4) | 55 | 0.03 |

